# Amplicon sequence variants artificially split bacterial genomes into separate clusters

**DOI:** 10.1101/2021.02.26.433139

**Authors:** Patrick D. Schloss

## Abstract

Amplicon sequencing variants (ASVs) have been proposed as an alternative to operational taxonomic units (OTUs) for analyzing microbial communities. ASVs have grown in popularity, in part, because of a desire to reflect a more refined level of taxonomy since they do not cluster sequences based on a distance-based threshold. However, ASVs and the use of overly narrow thresholds to identify OTUs increase the risk of splitting a single genome into separate clusters. To assess this risk, I analyzed the intragenomic variation of 16S rRNA genes from the bacterial genomes represented in a *rrn* copy number database, which contained 20,427 genomes from 5,972 species. As the number of copies of the 16S rRNA gene increased in a genome, the number of ASVs also increased. There was an average of 0.58 ASVs per copy of the 16S rRNA gene for full length 16S rRNA genes. It was necessary to use a distance threshold of 5.25% to cluster full length ASVs from the same genome into a single OTU with 95% confidence for genomes with 7 copies of the 16S rRNA, such as *E. coli*. This research highlights the risk of splitting a single bacterial genome into separate clusters when ASVs are used to analyze 16S rRNA gene sequence data. Although there is also a risk of clustering ASVs from different species into the same OTU when using broad distance thresholds, those risks are of less concern than artificially splitting a genome into separate ASVs and OTUs.

## Importance

16S rRNA gene sequencing has engendered significant interest in studying microbial communities. There has been a tension between trying to classify 16S rRNA gene sequences to increasingly lower taxonomic levels and the reality that those levels were defined using more sequence and physiological information than is available from a fragment of the 16S rRNA gene. Furthermore, naming of bacterial taxa reflects the biases of those who name them. One motivation for the recent push to adopt ASVs in place of OTUs in microbial community analyses is to allow researchers to perform their analyes at the finest possible level that reflects species-level taxonomy. The current research is significant because it quantifies the risk of artificially splitting bacterial genomes into separate clusters. Far from providing a better represenation of bacterial taxonomy and biology, the ASV approach can lead to conflicting inferences about the ecology of different ASVs from the same genome.

16S rRNA gene sequencing is a powerful technique for describing and comparing microbial communities (1). Efforts to link 16S rRNA gene sequences to taxonomic levels based on distance thresholds date to at least the 1990s. The distance-based threshold that was developed and is now widely used was based on DNA-DNA hybridization approaches that are not as precise as genome sequencing (2, 3). Instead, genome sequencing technologies have suggested that the widely used 3% distance threshold to operationally define bacterial taxa is too coarse (4–6). As an alternative to operational taxonomic units (OTUs), amplicon sequencing variants (ASVs) have been proposed as a way to adopt the thresholds suggested by genome sequencing to microbial community analysis using 16S rRNA gene sequences (7–10). Approaches for identifying ASVs do not cluster sequences based on a distance-based threshold (11). Proponents of ASVs are largely dissmissive of concerns that most bacterial genomes have more than one copy of the *rrn* operon and that those copies are not identical (12, 13). Yet, ASVs and using too fine a threshold to identify OTUs could split a single genome into multiple clusters. Conversely, using too broad of a threshold to define OTUs could cluster together multiple bacterial species into the same OTU. An example of both is seen in the comparison of *Staphylococcus aureus* (NCTC 8325) and *S. epidermidis* (ATCC 12228) where each genome has 5 copies of the 16S rRNA gene. Each of the 10 copies of the 16S rRNA gene in these two genomes is distinct and represent 10 ASVs. Conversely, if the copies were clustered using a 3% distance threshold, then all 10 ASVs would cluster into the same OTU. The goal of this study was to quantify the tradeoff of splitting a single genome into multiple clusters and the risk of clustering different bacterial species into the same cluster when using ASVs and various OTU definitions.

To investigate the variation in the number of copies of the 16S rRNA gene per genome and the intragenomic variation among copies of the 16S rRNA gene, I obtained 16S rRNA sequences from the *rrn* copy number database (*rrn*DB)(14). Among the 5,972 species represented in the *rrn*DB there were 20,427 genomes. The median *rrn* copy number per species ranged between 1 (e.g., *Mycobacterium tuberculosis*) and 19 (*Metabacillus litoralis*). As the *rrn* copy number for a genome increased, the number of variants of the 16S rRNA gene in each genome also increased. On average, there were 0.58 variants per copy of the full length 16S rRNA gene and an average of 0.32, 0.25, and 0.27 variants when considering the V3-V4, V4, and V4-V5 regions of the gene, respectively. Although a species tended to have a consistent number of 16S rRNA gene copies per genome, the number of total variants increased with the number of genomes that were sampled (Figure S1). For example, the 271 genome accessions of *Mycobacterium tuberculosis* in the *rrn*DB each had 1 copy of the gene per genome. However, across those accessions, there were 17 versions of the gene. An *E. coli* genome typically had 7 copies of the 16S rRNA gene with a median of 5 distinct full length ASVs per genome (intraquartile range between 3 and 6). Across the 1,390 *E. coli* genomes in the *rrn*DB, there were 1,402 versions of the gene. These observations highlight the risk of selecting a threshold for defining clusters that is too narrow because it is possible to split a single genome into multiple clusters.

A method to avoid splitting a single genome into multiple clusters is to cluster 16S rRNA gene sequences together based on their distances between each other. Therefore, I assessed the impact of the distance threshold used to define clusters of 16S rRNA genes on the propensity to split a genome into separate clusters. To control for uneven representation of genomes across species, I randomly selected one genome from each species and repeated each randomization 100 times. I observed that as the *rrn* copy number increased, the distance threshold required to reduce the ASVs in each genome to a single OTU increased (Figure 1). Among species with 7 copies of the *rrn* operon (e.g., *E. coli*), a distance threshold of 5.25% was required to reduce full length ASVs into a single OTU for 95% of the species. Similarly, thresholds of 5.25, 2.50, and 3.75% were required for the V3-V4, V4, and V4-V5 regions, respectively. But, if a 3% distance threshold was used, then ASVs from genomes containing fewer than 6, 6, 8, and 6 copies of the *rrn* operon would reliably be clustered into a single OTU for ASVs from the V1-V9, V3-V4, V4, and V4-V5 regions, respectively. Consequently, these results demonstrate that broad thresholds must be used to avoid splitting different operons from the same genome into separate clusters.

**Figure 1.**
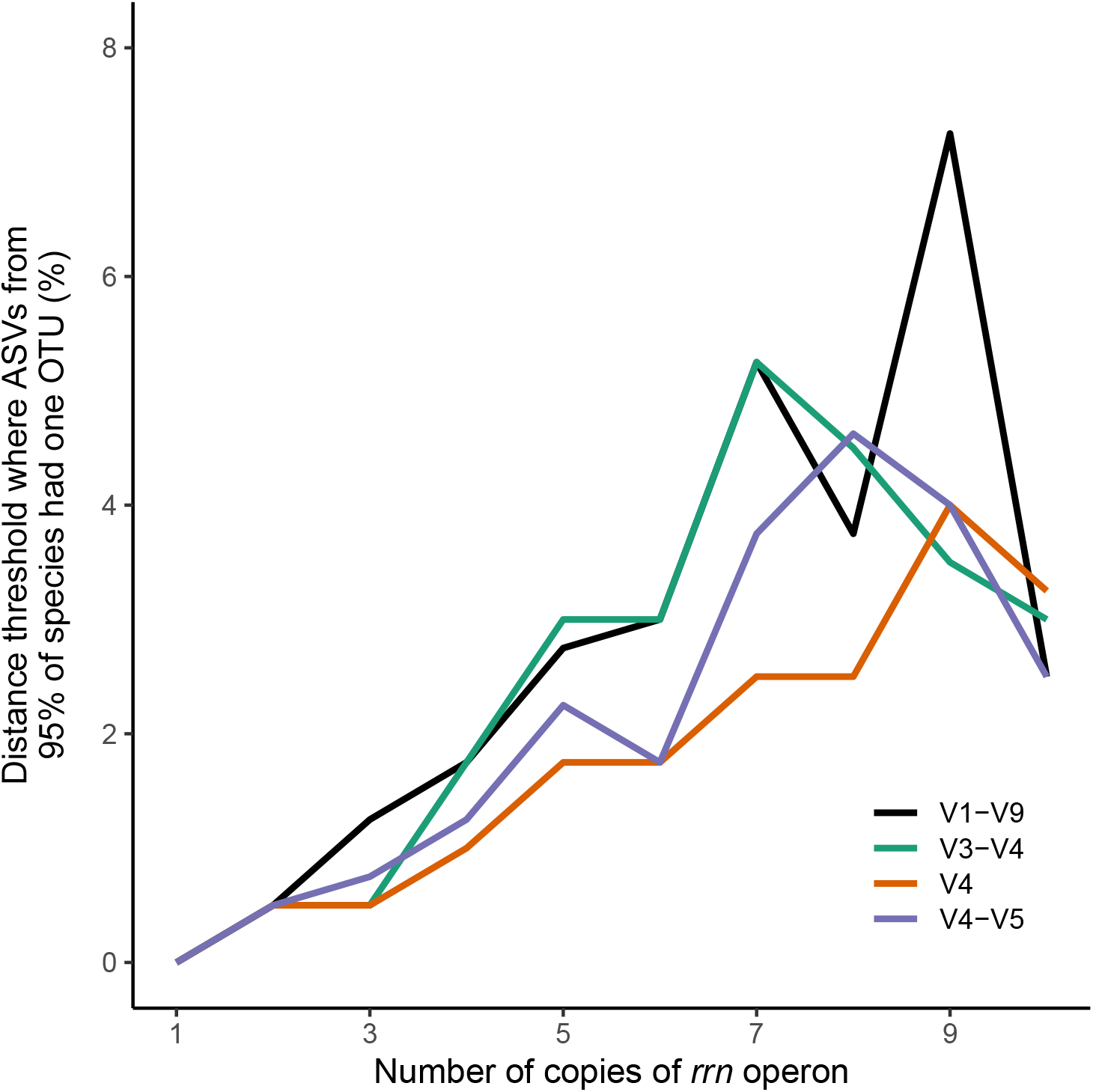
The distance threshold required to prevent the splitting of genomes into multiple OTUs increased as the number of *rrn* operons in the genome increased. Each line represents the median distance threshold for each region of the 16S rRNA gene that is required for 95% of the genomes with the indicated numbrer of *rrn* operons to cluster their ASVs to a single OTU. The median distance threshold was calculated across 100 randomizations in which one genome was sampled from each species. Only those number of *rrn* operons that were found in more than 100 species are included.

At broad thresholds, 16S rRNA gene sequences from multiple species could be clustered into the same ASV or OTU. I again randomly selected one genome from each species to control for uneven representation of genomes across species and for this analysis I measured the percentage of ASVs and OTUs that contained 16S rRNA gene sequences from multiple species (Figure 2). Without using distance-based thresholds, 4.1% of the ASVs contained sequences from multiple species when considering full length sequences and 10.9, 16.2, and 13.1% when considering the V3-V4, V4, and V4-V5 regions, respectively. At the commonly used 3% threshold for defining OTUs, 27.4% of the OTUs contained 16S rRNA gene sequences from multiple species when considering full length sequences and 31.7, 34.3, and 34.8% when considering the V3-V4, V4, and V4-V5 regions, respectively. Considering that species designations are inconsistently applied and reflect multiple human-imposed biases, the risk of splitting a genome into multiple OTUs is more problematic than clustering species together. Therefore, larger thresholds are advisable.

**Figure 2.**
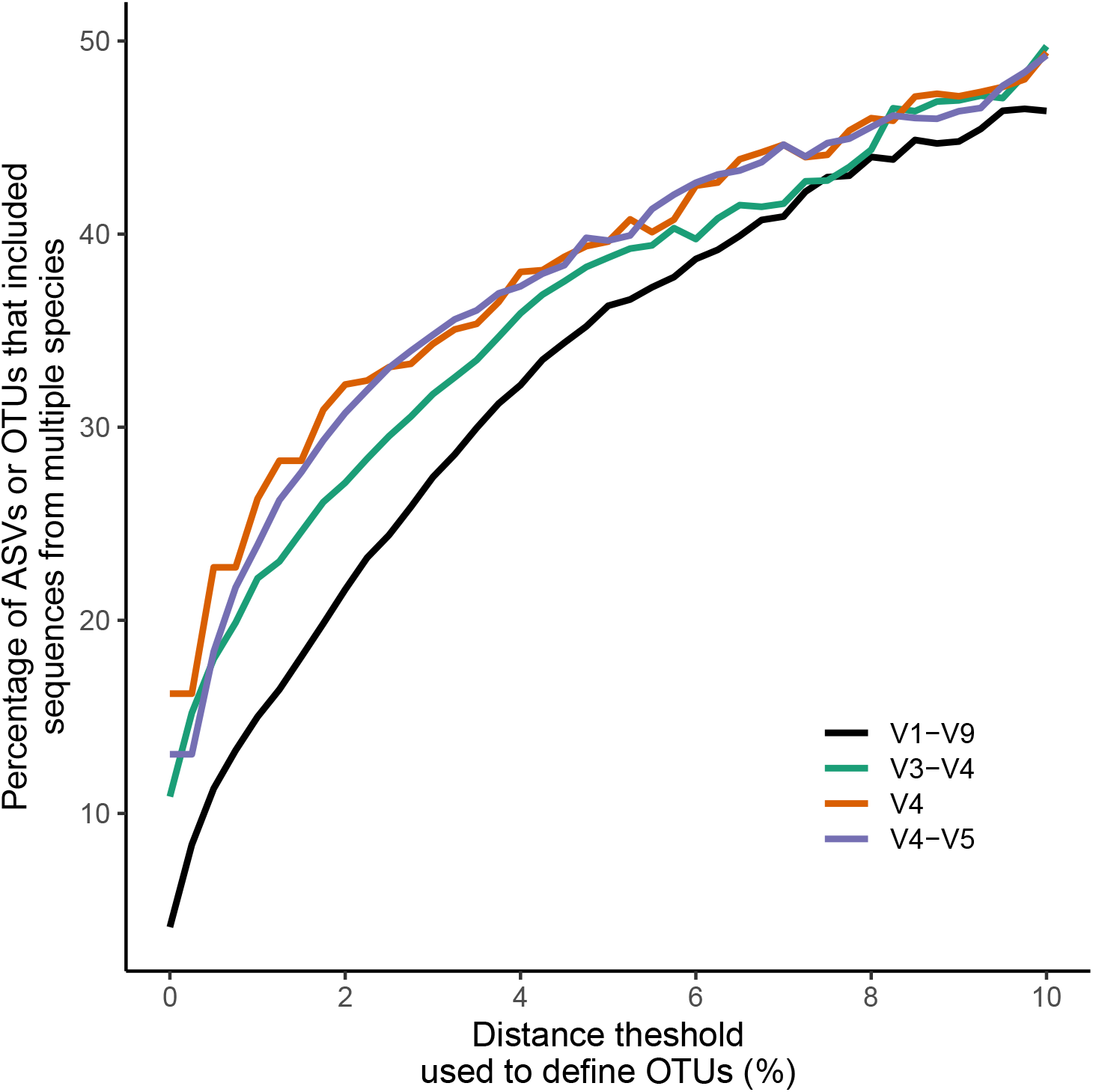
As the distance threshold used to define an OTU increased, the percentage of ASVs and OTUs representing multiple species increased. These data represent the median fractions for both measurements across 100 randomizations. In each randomization, one genome was sampled from each species.

The results of this analysis demonstrate that there is a significant risk of splitting a single genome into multiple clusters if using ASVs or too fine of a threshold to define OTUs. An ongoing problem for amplicon-based studies is defining a meaningful taxonomic unit (11, 15, 16). Since there is no consensus for a biologicaly definition of a bacterial species (17–19), microbiologists must accept that how bacterial species are named is biased and that taxonomic rules are not applied in a consistent manner (e.g., (19, 20)). This makes it impossible to fit a distance threshold to define an OTU definition that matches a set of species names (21). Furthermore, the 16S rRNA gene does not evolve at the same rate across all bacterial lineages (15), which limits the biological interpretation of a common OTU definition. A distance-based definition of a taxonomic unit based on 16S rRNA gene or full genome sequences is, at best, operational and not grounded in biological theory (15, 22–24). There is general agreement in bacterial systematics that to classify an organism to a bacterial species, phenotypic and genome sequence data are needed (17–20). A short sequence from a bacterial genome simply cannot differentiate between species. Moreover, it is difficult to defend a clustering threshold that would split a single genome into multiple taxonomic units. It is not biologically plausible to entertain the possibility that different *rrn* operons from the same genome would have different ecologies. Although there are multiple reasons that proponents favor ASVs, the significant risk of artificially splitting genomes into separate clusters is too high to warrant their use.

## Materials and Methods

### (i) Data availability

The 16S rRNA gene sequences used in this study were obtained from the *rrn*DB (https://rrndb.umms.med.umich.edu; version 5.7, released January 18, 2021) (14). At the time of submission, this was the most current version of the database. The *rrn*DB obtained the curated 16S rRNA gene sequences from the KEGG database, which ultimately obtained them from NCBI’s non-redundant RefSeq database. The *rrn*DB provided downloadable versions of the sequences with their taxonomy as determined using the naive Bayesian classifier trained on the RDP reference taxonomy. For some genomes this resulted in multiple classifications since a genome’s 16S rRNA gene sequences were not identical. Instead, I mapped the RefSeq accession number for each genome in the database to obtain a single taxonomy for each genome. Because strain names were not consistently given to genomes across bacterial species, I disregarded the strain level designations.

### (ii) Definition of regions within the 16S rRNA gene

The full length 16S rRNA gene sequences were aligned to a SILVA reference alignment of the 16S rRNA gene (v. 138) using the mothur software package (v. 1.44.2) (25, 26). Regions of the 16S rRNA gene were selected because of their use in the microbial ecology literature. Full length sequences corresponded to *E. coli* str. K-12 substr. MG1655 (NC_000913) positions 28 through 1491, V4 to positions 534 through 786, V3-V4 to positions 358 through 786, and V4-V5 to positions 534 through 908. The positions between these coordinates reflect the fragments that would be amplified using commonly used PCR primers.

### (iii) Clustering sequences into OTUs

Pairwise distances between sequences were calculated using the dist.seqs command from mothur. The OptiClust algorithm, as implemented in mothur, was used to assign 16S rRNA gene sequences to OTUs (27). Distance thresholds between 0.25 and 10.00% in 0.25 percentage point increments were used to assign sequences to OTUs.

### (iv) Controlling for uneven sampling of genomes by species

Because of the uneven distribution of genome sequences across species I randomly selected one genome from each species for the analysis of splitting genomes and clustering ASVs from different species (Figures 1 and 2). The random selection was repeated 100 times. Analyses based on this randomization reported the median of the 100 randomizations. The intraquartile range between randomizations was less than 0.0024. Because the range was so small, the confidence intervals were more narrow than the thickness of the lines in Figures 1 and 2 and were not included.

### (v) Reproducible data analysis

The code to perform the analysis in this manuscript and its history are available as a git-based version control repository on GitHub (https://github.com/SchlossLab/Schloss_rrnAnalysis_mSphere_2021). The analysis can be regenerated using a GNU Make-based workflow that made use of built-in bash tools (v. 3.2.57), mothur (v. 1.44.2), and R (v. 4.0.4). Within R, I used the tidyverse (v. 1.3.0), data.table (v. 1.13.2), Rcpp (v. 1.0.5), furrr (v. 0.2.1), here (v. 1.0.1) and rmarkdown (v. 2.5) packages. The conception and development of this analysis is available as a playlist on the Riffomonas YouTube channel (https://youtube.com/playlist?list=PLmNrK_nkqBpL7m_tyWdQgdyurerttCsPY).

### (vi) Note on usage of ASV, OTU, and cluster

I used “ASV” to denote the cluster of true 16S rRNA gene sequences that were identical to each other and “OTU” to denote the product of distance-based clustering of sequences. Although ASVs do represent a type of operational defition of a taxonomic unit and can be thought of as an OTU formed using a distance of zero, proponents of the ASV approach prefer to avoid the term OTU given the long history of OTUs being formed by distance-based clustering (https://github.com/benjjneb/dada2/issues/62; accessed 2021-02-26). For this reason, when an ASV split a genome into different units, those units were called clusters rather than OTUs.

## Acknowledgements

I am grateful to Robert Hein and Thomas Schmidt, who maintain the *rrn*DB, for their help in understanding the curation of the database and for making the 16S rRNA gene sequences and related metadata publicly available. I am also grateful to community members who watched the serialized version of this analysis on YouTube and provided suggestions and questions over the course of the development of this project. This work was supported, in part, through grants from the NIH (P30DK034933, U01AI124255, and R01CA215574).

---

**Figure S1.**
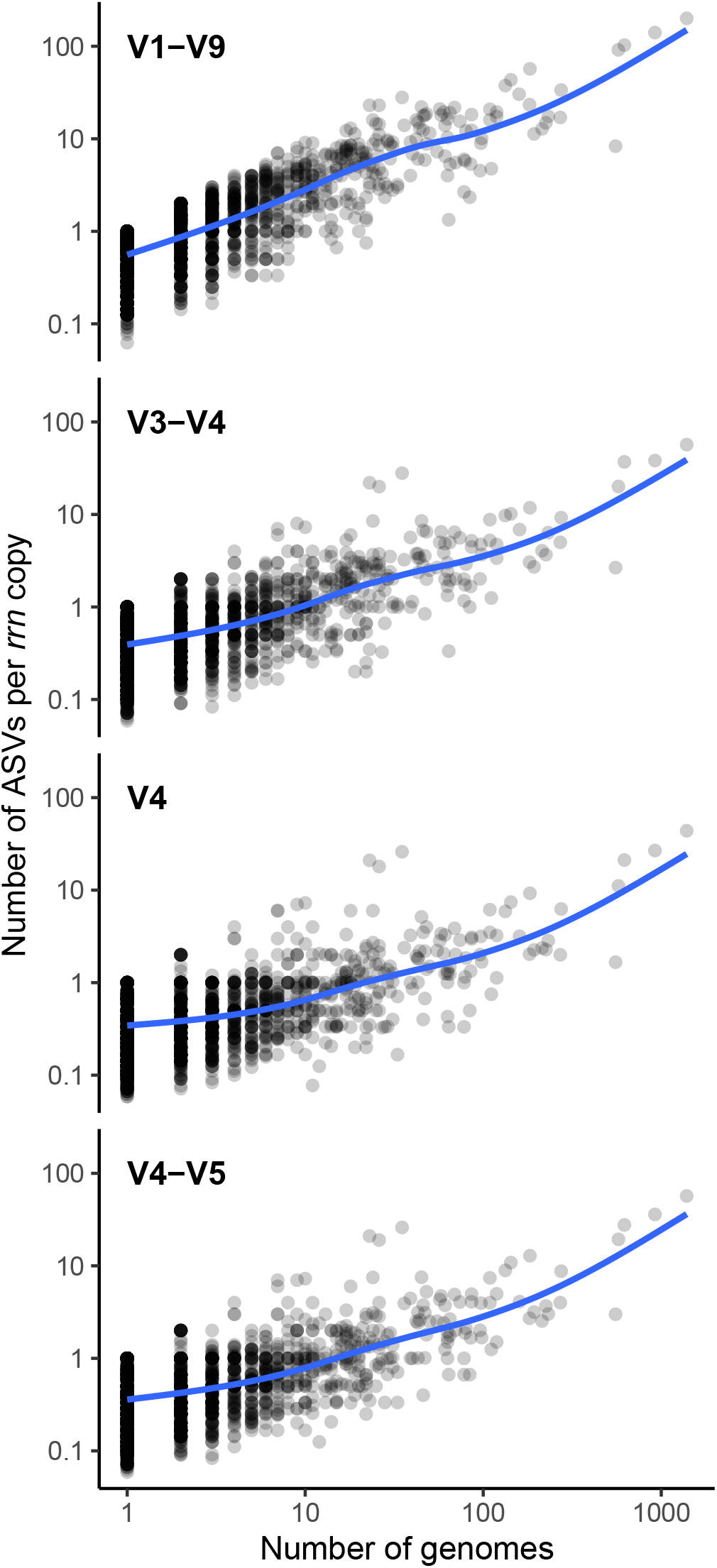
The ratio of number of distinct ASVs per copy of the *rrn* operon increased for a species as the number of genomes in the *rrn*DB for that species increased. Each point represents a different species and was shaded to be 80% transparent so that when points overlap they become darker. The blue line represents a smoothed fit through the data. Both axes use a logarithmic scale (base 10).

